# A Bioinformatic Study of the Distribution of Mn Oxidation Proteins in Sequenced Bacterial Genomes

**DOI:** 10.1101/2022.11.10.515945

**Authors:** M. Zakaria Kurdi, Jacob Olichney, Kati Geszvain

## Abstract

1.

**Background:** While many species of bacteria have been identified that can convert soluble, reduced manganese (Mn^+2^) into insoluble, oxidized Mn^+4^ oxides, the mechanisms these bacteria employ and their distribution throughout the bacterial domain are less well understood. One of the best characterized MnOB is the gamma-proteobacterium *Pseudomonas putida* GB-1, which uses three distinct proteins (PpMnxG, McoA and MopA) to oxidize Mn^+2^. The best characterized Mn oxidase enzyme is the MnxG homolog of *Bacillus* sp. PL-12 (BaMnxG), which appears to be the only Mn oxidase in this species. MofA, found in *Leptothrix discophora* sp SS-1 is an additional putative Mn oxidase.

**Results:** By querying publicly available databases of bacterial genome sequences for homologs to these Mn oxidase proteins, it was possible to determine the distribution of the proteins within bacteria. The overwhelming majority of homologs were found in just three phyla: proteobacteria, actinobacteria and firmicutes. These data do not preclude the possibility of novel Mn oxidase mechanisms in other as yet uncharacterized groups of bacteria. Each of the homologs had a statistically significant probability of being present as the solo Mn oxidase in a genome. When genomes did have more than one oxidase, they were present in the same combinations as in *P. putida* GB-1.

**Conclusions:** These results do not support the initial hypothesis that multiple enzymes are required to complete the two-electron oxidation of Mn^+2^ to Mn^+4^. Alternatively, the various Mn oxidase enzymes may be optimized to function under different environmental conditions; organisms like *P. putida* GB-1 may need to oxidize Mn at different temperatures, nutritional states or oxygen conditions.

## 2. Background

Manganese, a common transition metal in the Earth’s crust, cycles between the reduced, soluble form Mn^+2^ and oxidized Mn^+4^ forms which are insoluble, highly reactive and readily adsorb metals and other compounds in their environment (1). While oxidation of Mn can occur abiotically, the rate of this reaction is increased up to five orders of magnitude by bacteria and fungi (2) and the biogenic Mn oxides that result are strong oxidants currently being investigated for their use in bioremediation of contaminated wastewater (2–4). Understanding the organisms that oxidize manganese and the genes they use to do so will be essential for this application.

Mn-oxidizing bacteria (MnOB) are a diverse group found in many phyla including both Gram-positive and Gram-negative bacteria (5). Within the Gram-negative organisms, several classes within the phylum proteobacterium are represented, including alpha-, beta- and gamma-proteobacteria (6,7). Other Gram-negative MnOB fall in the genus *Flavobacterium* within the phylum Bacteroidetes (8). The Gram-positive MnOB include many phylogenetically distinct species of *Bacillus* in the phylum Firmicutes (9– 11), and some species in the genus *Arthrobacter*, in the phylum Actinobacteria (12).

While much of the work in the field of MnOB has focused on identifying the bacteria that carry out this activity in a given environment, some work has also been done to identify the enzymes used to oxidize Mn. The best characterized Mn oxidase enzyme is the Mnx complex from *Bacillus* sp. PL-12 (13–16). This complex is composed of three proteins, MnxE, F and G (14). PpMnxG, the largest of the three proteins, belongs to the multicopper oxidase (MCO) protein family and is likely the catalytic component of the complex (14,17). Proteins belonging to the peroxidase cyclooxygenase family have also been identified as Mn oxidases (18,19). Peroxidase cyclooxygenases, also referred to as animal heme peroxidases (AHP) are characterized by a heme-binding domain and are found in animals, fungi, plants and bacteria (20).

The well characterized MnOB *Pseudomonas putida* GB-1 has been shown to use three separate Mn oxidase enzymes (21,22). Two of the enzymes - McoA and MnxG - belong to the MCO family (22). The third Mn oxidase gene encodes the AHP MopA (21). It is not known how common it is for a bacterium to encode multiple Mn oxidase enzymes nor is it known what purpose it serves to have multiple oxidase enzymes. However, the genome of the MnOB *Leptothrix discophora* sp SS-1 is also predicted to encode multiple Mn oxidases (23).

In addition to the MnxG Mn oxidases found in *Bacillus* sp. SG-1 and *P. putida* GB-1, other MCO-type Mn oxidases include MofA, found in *Leptothrix discophora* sp. SS-1 (24) and MoxA from *Pedomicrobium* sp. ACM 3067 (25) as well as CotA, CopA, MokA and CueO in *Bacillus* sp. (26,27), *Brevibacillus panacihumi* sp. MK-8 (28), *Lysinibacillus* sp. MK-1 (29) and *Escherichia coli* (30), respectively. The MCO Mn oxidases vary in their size, and the number of predicted Cu-binding domains (Table 1). In a pairwise BLAST comparison of amino acid sequence (Table 3), all of the MCO Mn oxidases have a low level of homology to one another, but they also have similar levels of homology to CumA, a non-Mn-oxidizing MCO (31). The closest homology is between *Bacillus* sp. MnxG (BaMnxG) and *P. putida* GB-1 MnxG (PpMnxG) with 25% identity over 99% coverage.

**Table 1.**
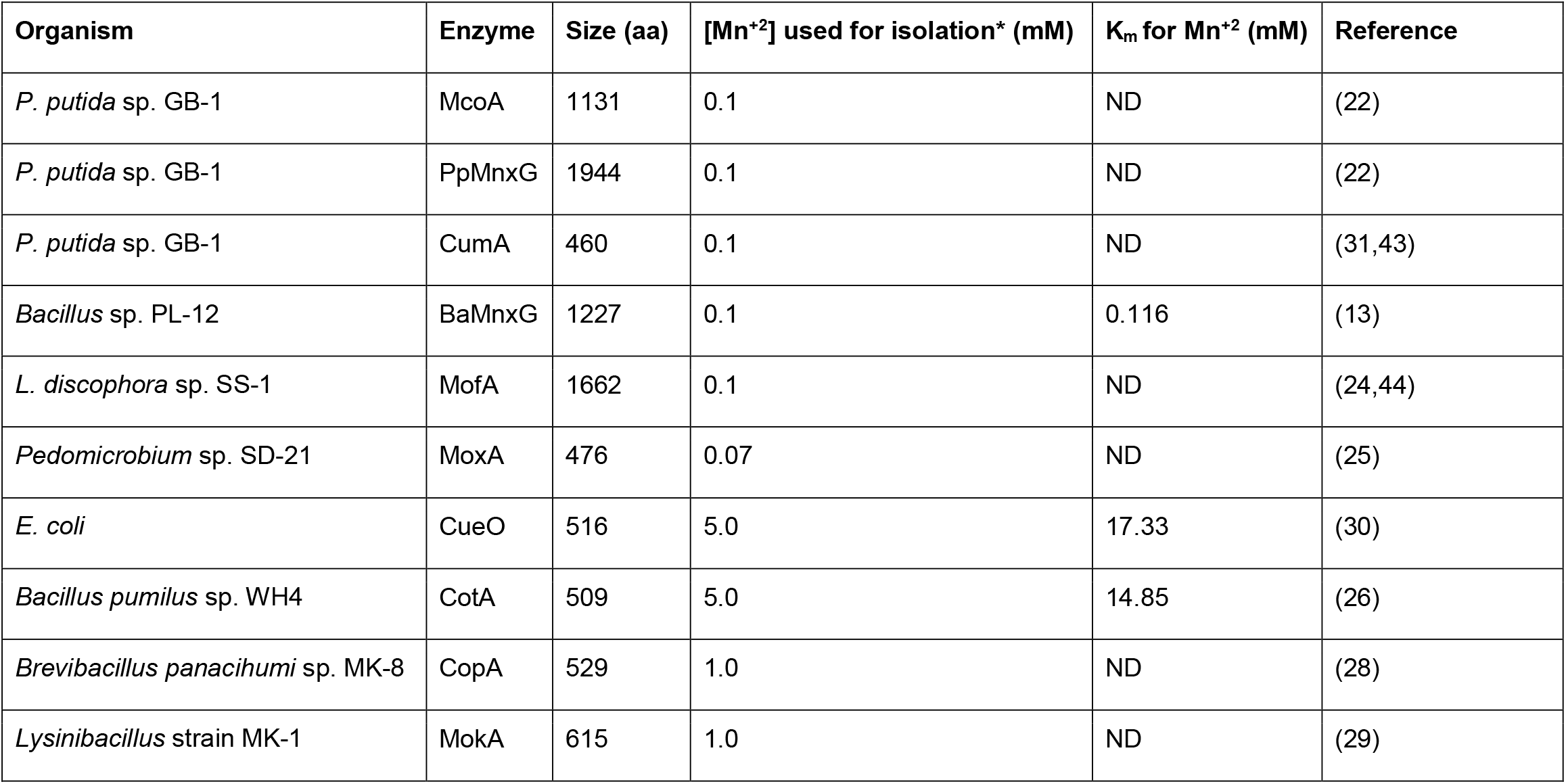
MCO Mn oxidases

| Organism | Enzyme | Size (aa) | [Mn<sup>+2</sup>] used for isolation* (mM) | K<sub>m</sub> for Mn<sup>+2</sup> (mM) | Reference |
| --- | --- | --- | --- | --- | --- |
| *P. putida* sp. GB-1 | McoA | 1131 | 0.1 | ND | (22) |
| *P. putida* sp. GB-1 | PpMnxG | 1944 | 0.1 | ND | (22) |
| *P. putida* sp. GB-1 | CumA | 460 | 0.1 | ND | (31,43) |
| *Bacillus* sp. PL-12 | BaMnxG | 1227 | 0.1 | 0.116 | (13) |
| *L. discophora* sp. SS-1 | MofA | 1662 | 0.1 | ND | (24,44) |
| *Pedomicrobium* sp. SD-21 | MoxA | 476 | 0.07 | ND | (25) |
| *E. coli* | CueO | 516 | 5.0 | 17.33 | (30) |
| *Bacillus pumilus* sp. WH4 | CotA | 509 | 5.0 | 14.85 | (26) |
| *Brevibacillus panacihumi* sp. MK-8 | CopA | 529 | 1.0 | ND | (28) |
| *Lysinibacillus* strain MK-1 | MokA | 615 | 1.0 | ND | (29) |

**Table 2.** AHP Mn oxidases

| Organism | Enzyme | Size (aa) | [Mn<sup>+2</sup>] used for isolation* (mM) | # heme domains | Km for Mn<sup>+2</sup> (mM) | Reference |
| --- | --- | --- | --- | --- | --- | --- |
| *A. maganoxydans* sp<br>SI85-9A1 | MopA | 3279 | 0.1 | 2 | ND | (18) |
| *Erythrobacter* sp. SD-21 | MopA | 2137 | 0.1 | 1 | 0.154 (heme domain) | (18,32,33) |
| *P. putida* GB-1 | MopA | 3608 | 0.1 | 2 | ND | (21) |
| *Roseobacter* sp. AzwK-3b | AhpA | 563 | 0.1 | 0 | ND | (19,45) |
|  | AhpB | 1706 |  | 1 | ND |  |
|  | AhpC | 1277 |  | 1 | ND |  |
|  | AhpL | 3578 |  | 2 | ND |  |

**Table 3.** Homology within the MCO class of Mn oxidases (% identity, % coverage). The non-Mn-oxidizing MCO CumA is included for comparison. NSS = No Significant Similarity, Red = >20% identical and >50% coverage

|  | BaMnxG | MofA | MoxA | McoA | PpMnxG | CueO | CotA | CopA | MokA | CumA |
| --- | --- | --- | --- | --- | --- | --- | --- | --- | --- | --- |
| BaMnxG |  | NSS | 26, 5 | NSS | 24, 99 | NSS | NSS | 23, 26 | NSS | NSS |
| MofA |  |  | 25, 5 | 23, 20 | NSS | 28, 11 | 28, 37 | 29, 3 | 24, 34 | 34, 11 |
| MoxA |  |  |  | NSS | NSS | 39, 35 | 28, 24 | 28, 54 | 28, 20 | 33, 45 |
| McoA |  |  |  |  | NSS | 32, 18 | 24, 37 | 30, 15 | 27, 30 | 28, 16 |
| PpMnxG |  |  |  |  |  | NSS | NSS | 29, 11 | NSS | NSS |
| CueO |  |  |  |  |  |  | 25, 92 | 25, 86 | 25, 89 | 25, 59 |
| CotA |  |  |  |  |  |  |  | 24, 89 | 52, 99 | 21, 91 |
| CopA |  |  |  |  |  |  |  |  | 22, 80 | 28, 86 |
| MokA |  |  |  |  |  |  |  |  |  | 21, 78 |
| CumA |  |  |  |  |  |  |  |  |  |  |

Three MCO Mn oxidase enzymes have been biochemically characterized thus far. The *Bacillus* sp. PL-12 Mnx complex has a K_m_ for Mn^+2^ of 0.116 mM. This is similar to the K_m_ for the *Erythrobacter* SD-21 MopA heme domain (32,33) (Table 2). However, these K_m_ values are an order of magnitude lower than that for CueO and CotA, the Mn oxidase enzymes of *E. coli* and *Bacillus pumilus* sp. WH4 respectively. Possibly relevant, these later two organisms were isolated from the environment using a much higher concentration of Mn^+2^ than was used for *Bacillus* sp. PL-12 and *Erythrobacter* sp. SD-21(Table 1).

Biochemical purification of the Mn oxidase activity from *Aurantimonas manganoxydans* SI85-9A1 and *Erythrobacter* sp. SD-21 identified an AHP subsequently named MopA, a large protein predicted to have two heme-binding domains in *A. manganoxydans* sp SI85-9A1 and one in *Erythrobacter* sp. SD-21 (18). Heterologous expression of the heme-binding domain from the *Erythrobacter* sp SD-21 ErMopA in *E. coli* resulted in a partially-purified protein capable of oxidizing Mn^+2^ to Mn^+3^ (32,33). Four AHPs have also been implicated in Mn oxidation by the marine bacterium *Roseobacter* sp AzwK-3b (19).

The AHP Mn oxidase enzymes vary in size from the 563 amino acid AhpA to the 3608 amino acid AhpL. Except for AhpA, each has either one or two predicted heme-binding domains (Table 2). BLASTP pairwise comparison of their amino acid sequences shows a substantial amount of similarity among these enzymes (Table 4). This is likely due to the size of the conserved heme-binding domains found within each protein.

**Table 4.** Homology between AHP class of Mn oxidases (% identity, % coverage) NSS = No Significant Similarity, Red = >20% identical and >50% coverage

|  | **PpMopA** | **AmMopA** | **ErMopA** | **AhpA** | **AhpB** | **AhpC** | **AhpL** |
| --- | --- | --- | --- | --- | --- | --- | --- |
| **PpMopA** |  | 52, 94 | 42, 94 | 46, 21 | 47, 76 | 30, 51 | 48, 84 |
| **AmMopA** |  |  | 57, 83 | 50, 23 | 53, 78 | 52, 61 | 55, 93 |
| **ErMopA** |  |  |  | 47, 24 | 48, 73 | 31, 51 | 49, 89 |
| **AhpA** |  |  |  |  | NSS | NSS | 100, 93 |
| **AhpB** |  |  |  |  |  | 27, 52 | 99, 99 |
| **AhpC** |  |  |  |  |  |  | 100, 100 |
| **AhpL** |  |  |  |  |  |  |  |

The identification of multiple Mn oxidase enzymes and the availability of a large number of sequenced bacterial genomes makes it possible to address these questions regarding the distribution of Mn oxidase proteins. In the first section of the paper, we will review the literature to determine which Mn oxidation proteins to use in our analysis. We will then use the amino acid sequence of these proteins to query publicly available databases of protein sequences to identify putative homologs and generate a database of bacterial genomes that include at least one putative Mn oxidase homolog. Our results will show that Mn oxidase proteins are predominantly found in just three phyla and that these three phyla differ in which Mn oxidase proteins are present. Furthermore, we will show that Mn oxidase proteins are commonly found in isolation, suggesting that *P. putida* GB-1 is unusual in having three Mn oxidase proteins in its genome.

## 3. Results

### Selection of Mn oxidase proteins for further analysis

A subset of the previously identified Mn oxidase proteins was selected for further analysis. As the best characterized Mn oxidase, MnxG from the *Bacillus* sp. PL-12 Mnx complex (BaMnxG) and its homolog MnxG from *P. putida* GB-1 (PpMnxG, 25% identical, 99% coverage) were both chosen. Genetic evidence also supports the identification of McoA from *P. putida* GB-1 and MofA from *L. discophora* SS-1 as Mn oxidase enzymes. The putative Mn oxidases CueO and CotA have been shown in vitro to oxidize Mn; however, the K_m_ for these enzymes is 10-20-fold higher than BaMnxG or ErMopA-heme (Tables 1 and 2), suggesting that these enzymes biochemically are quite different from MnxG and MopA. Similarly, CopA and MokA were identified as Mn oxidases after screening for MnOB at a Mn^+2^ concentration that is an order of magnitude higher than that used to identify other MnOB (Tables 1 and 2). These four enzymes are also more similar to each other and the non-oxidizing MCO CumA than to the other Mn oxidases (Table 3). Therefore, CopA, CotA, CueO and MokA were excluded from this study.

*Pedomicrobium* sp. ACM 3067 was isolated as an MnOB using a relatively low concentration of Mn^+2^ [66 µM, (25)]. However, the MCO tentatively identified as its Mn oxidase enzyme, MoxA, is similar in size to CumA, CueO, CotA, MokA and CopA. The genetic identification of MoxA as a Mn oxidase may also not be reliable (18). In preliminary experiments addressing the distribution of the Mn oxidase proteins in publicly available databases, MoxA homologs were extremely common. For these reasons, we decided that MoxA may not specifically oxidize Mn^+2^ and it was excluded from our studies.

We also included Mn oxidase proteins in the AHP class. The identification of the AHP MopA in *Erythrobacter* sp. SD-21 as a Mn oxidase enzyme is supported by both genetic and biochemical evidence and has a Km for Mn similar to that of BaMnxG. A similar AHP, also named MopA, in *P. putida* GB-1 has been shown genetically to be involved in Mn oxidation. Because of the relatively high conservation among the seven AHP Mn oxidase enzymes (Table 4), we chose to only screen for homologs to the heme-binding domain of *P. putida* GB-1 protein MopA.

### Distribution of Putative Mn Oxidase Enzymes

We screened the NCBI nr and IMG databases (34,35) for homologs to the Mn oxidase proteins BaMnxG (Phylum Firmicutes, Class Bacillus), PpMnxG (Proteobacteria, gammaproteobacteria), McoA (Proteobacteria, gammaproteobacteria), MofA (Proteobacteria, betaproteobacteria), and MopA (Proteobacteria, gammaproteobacteria), and identified 2197 putative homologs in 1695 genomes (data not shown). MopA homologs were identified most frequently (1227/2197) while the other four enzymes were less than one third as frequent (Table 5). MofA and BaMnxG were the least frequently identified homologs (148 and 136, respectively, Table 5).

**Table 5.**
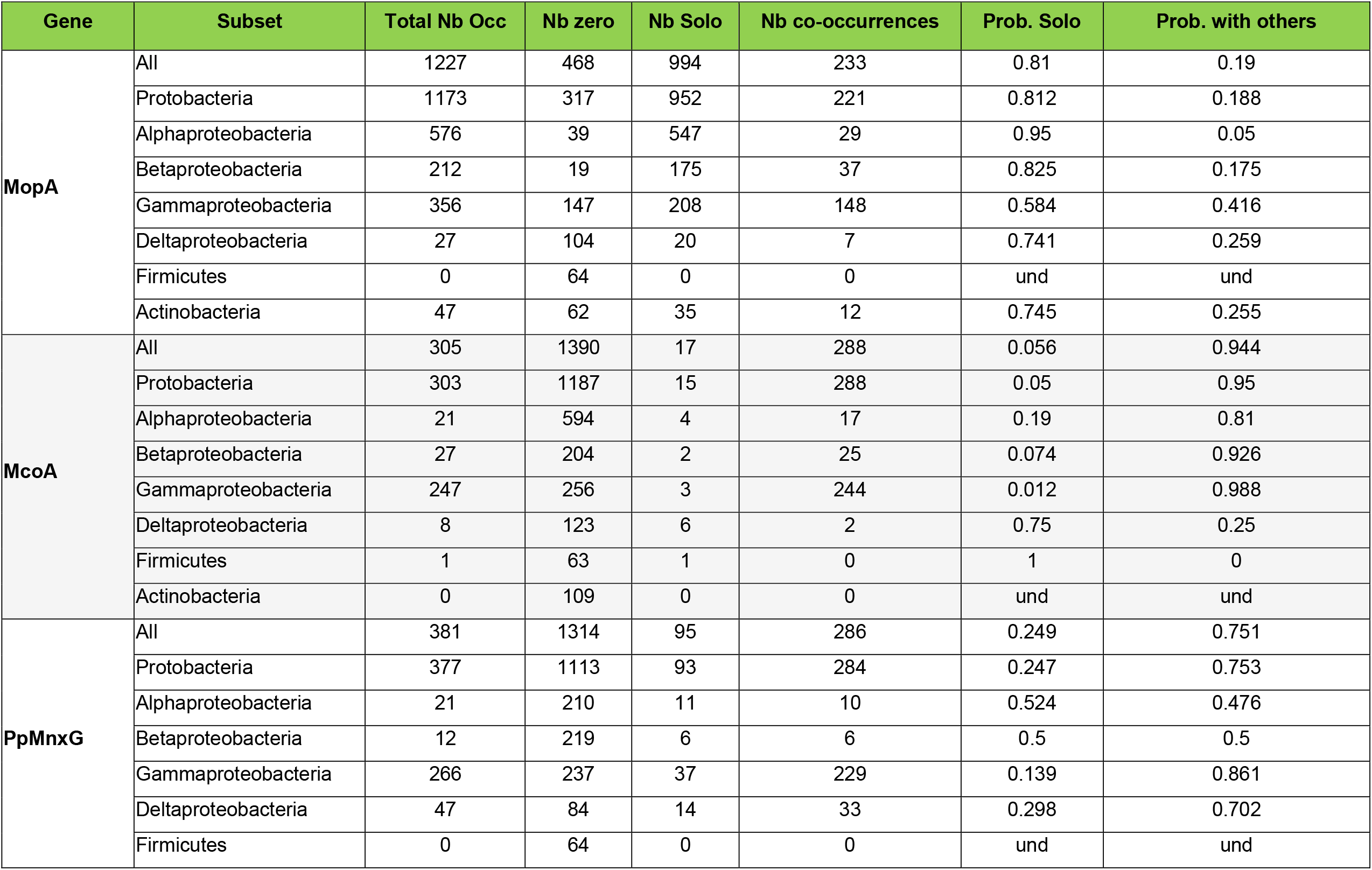
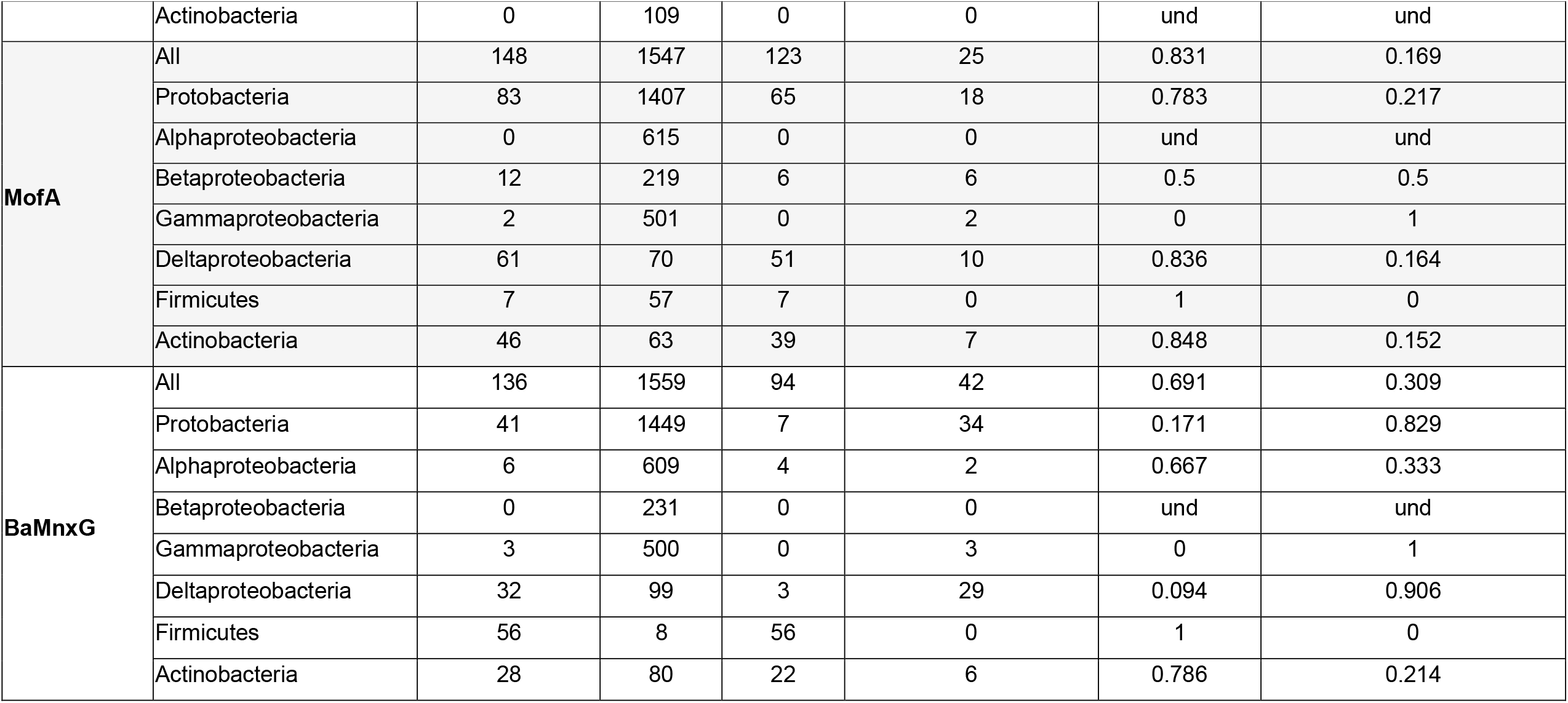
Distribution of manganese oxidation genes within the dataset

| Gene | Subset | Total Nb Occ | Nb zero | Nb Solo | Nb co-occurrences | Prob. Solo | Prob. with others |
| --- | --- | --- | --- | --- | --- | --- | --- |
| **MopA** | All | 1227 | 468 | 994 | 233 | 0.81 | 0.19 |
|  | Protobacteria | 1173 | 317 | 952 | 221 | 0.812 | 0.188 |
|  | Alphaproteobacteria | 576 | 39 | 547 | 29 | 0.95 | 0.05 |
|  | Betaproteobacteria | 212 | 19 | 175 | 37 | 0.825 | 0.175 |
|  | Gammaproteobacteria | 356 | 147 | 208 | 148 | 0.584 | 0.416 |
|  | Deltaproteobacteria | 27 | 104 | 20 | 7 | 0.741 | 0.259 |
|  | Firmicutes | 0 | 64 | 0 | 0 | und | und |
|  | Actinobacteria | 47 | 62 | 35 | 12 | 0.745 | 0.255 |
| **McoA** | All | 305 | 1390 | 17 | 288 | 0.056 | 0.944 |
|  | Protobacteria | 303 | 1187 | 15 | 288 | 0.05 | 0.95 |
|  | Alphaproteobacteria | 21 | 594 | 4 | 17 | 0.19 | 0.81 |
|  | Betaproteobacteria | 27 | 204 | 2 | 25 | 0.074 | 0.926 |
|  | Gammaproteobacteria | 247 | 256 | 3 | 244 | 0.012 | 0.988 |
|  | Deltaproteobacteria | 8 | 123 | 6 | 2 | 0.75 | 0.25 |
|  | Firmicutes | 1 | 63 | 1 | 0 | 1 | 0 |
|  | Actinobacteria | 0 | 109 | 0 | 0 | und | und |
| **PpMnxG** | All | 381 | 1314 | 95 | 286 | 0.249 | 0.751 |
|  | Protobacteria | 377 | 1113 | 93 | 284 | 0.247 | 0.753 |
|  | Alphaproteobacteria | 21 | 210 | 11 | 10 | 0.524 | 0.476 |
|  | Betaproteobacteria | 12 | 219 | 6 | 6 | 0.5 | 0.5 |
|  | Gammaproteobacteria | 266 | 237 | 37 | 229 | 0.139 | 0.861 |
|  | Deltaproteobacteria | 47 | 84 | 14 | 33 | 0.298 | 0.702 |
|  | Firmicutes | 0 | 64 | 0 | 0 | und | und |
|  | Actinobacteria | 0 | 109 | 0 | 0 | und | und |
| MofA | All | 148 | 1547 | 123 | 25 | 0.831 | 0.169 |
|  | Protobacteria | 83 | 1407 | 65 | 18 | 0.783 | 0.217 |
|  | Alphaproteobacteria | 0 | 615 | 0 | 0 | und | und |
|  | Betaproteobacteria | 12 | 219 | 6 | 6 | 0.5 | 0.5 |
|  | Gammaproteobacteria | 2 | 501 | 0 | 2 | 0 | 1 |
|  | Deltaproteobacteria | 61 | 70 | 51 | 10 | 0.836 | 0.164 |
|  | Firmicutes | 7 | 57 | 7 | 0 | 1 | 0 |
|  | Actinobacteria | 46 | 63 | 39 | 7 | 0.848 | 0.152 |
| BaMnxG | All | 136 | 1559 | 94 | 42 | 0.691 | 0.309 |
|  | Protobacteria | 41 | 1449 | 7 | 34 | 0.171 | 0.829 |
|  | Alphaproteobacteria | 6 | 609 | 4 | 2 | 0.667 | 0.333 |
|  | Betaproteobacteria | 0 | 231 | 0 | 0 | und | und |
|  | Gammaproteobacteria | 3 | 500 | 0 | 3 | 0 | 1 |
|  | Deltaproteobacteria | 32 | 99 | 3 | 29 | 0.094 | 0.906 |
|  | Firmicutes | 56 | 8 | 56 | 0 | 1 | 0 |
|  | Actinobacteria | 28 | 80 | 22 | 6 | 0.786 | 0.214 |

Fourteen separate phyla included genomes encoding at least one Mn oxidase homolog (data not shown). However, three phyla are responsible for 98% of the Mn oxidase homologs; most of the homologs can be found in the phylum proteobacteria (1977), while the actinobacteria (122) and firmicutes (64) phyla also have substantial numbers of homologs (Table 6). The three phyla that have the most putative Mn oxidase homologs each exhibit unique patterns of Mn oxidase genes (Table 6). In the Firmicutes, 56 out of 64 Mn oxidase homologs belong to the Bacillus-type MnxG (BaMnxG) group, 7 belong to the MofA group and 1 to the McoA group. The Actinobacteria phylum has 46 MofA, 47 MopA and 29 BaMnxG homologs. The proteobacteria phylum has representatives of each of the 5 putative Mn oxidase enzymes. In the gamma-, alpha- and betaproteobacteria, MopA, McoA and PpMnxG dominate. In gammaproteobacteria, these three enzymes are found with roughly equivalent frequency. However, in the alpha- and betaproteobacteria MopA is by far the most common putative Mn oxidase homolog present. The deltaproteobacteria class is unusual in that it possesses a substantial number of homologs to BaMnxG and even more to MofA (Table 6).

**Table 6.** Distribution of Mn oxidation genes by taxon within the dataset

| **Taxon** | **BaMnxG** | **PpMnxG** | **McoA** | **MopA** | **MofA** | **Total** |
| --- | --- | --- | --- | --- | --- | --- |
| **Firmicutes** | 56 | 0 | 1 | 0 | 7 | 64 |
| **Actinobacteria** | 29 | 0 | 0 | 47 | 46 | 122 |
| **Proteobacteria** | 41 | 377 | 303 | 1173 | 83 | 1977 |
| **Alpha-** | 6 | 21 | 21 | 576 | 0 | 624 |
| **Beta-** | 0 | 12 | 27 | 212 | 12 | 263 |
| **Gamma-** | 3 | 266 | 247 | 356 | 2 | 874 |
| **Delta-** | 32 | 47 | 8 | 27 | 61 | 155 |

The Shannon diversity index reflects these differences in homolog distribution. As shown in table 7, Actinobacteria and Proteobacteria both have high diversity indices resulting from the relatively even distribution of multiple Mn oxidase proteins in these phyla. However, the Firmicutes have a much lower diversity score due to the predominance of BaMnxG in this group.

**Table 7.**
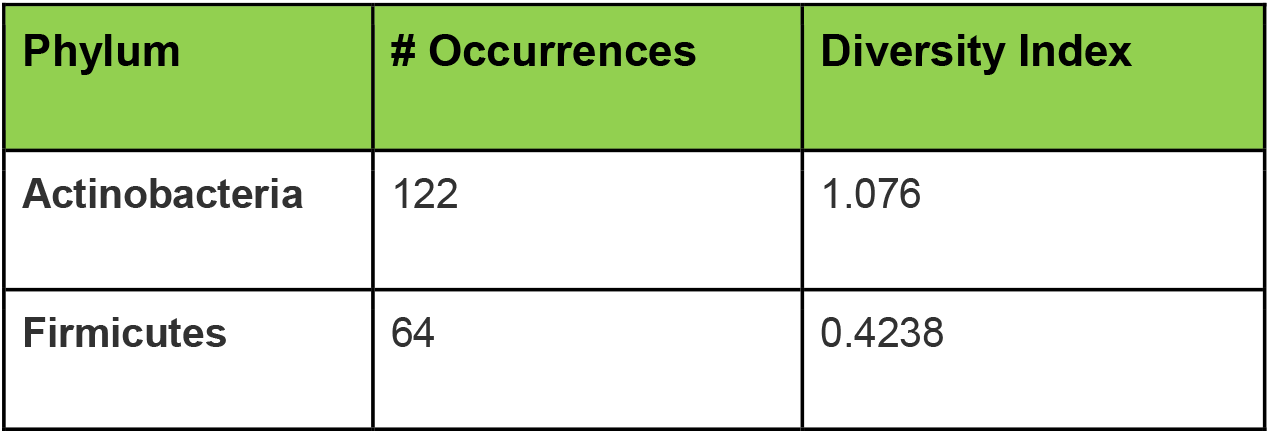
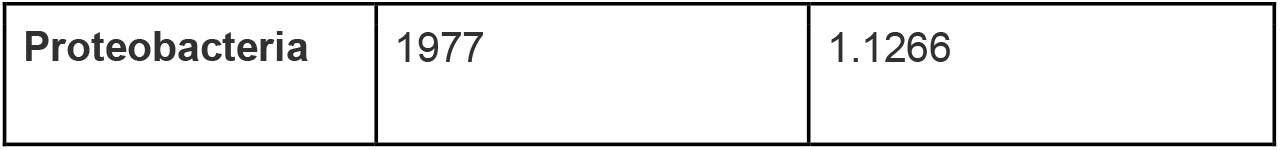
Shannon’s Diversity indices of the phyla

### Co-occurrence of Mn oxidase genes

*Pseudomonas putida* GB-1 has been shown to encode three separate Mn oxidase genes in its genome (21,22) and the genome of the MnOB *L. discophora* sp. SS-1 encodes multiple putative Mn oxidases (23). This raises the possibility that in some species Mn oxidation requires multiple enzymes working together. If this is true, we would predict that genomes would commonly have more than one Mn oxidase gene present. We addressed this question by determining the probability that each Mn oxidase protein was the only oxidase present (Prob. Solo, Table 5) vs the probability it co-occurred with another oxidase (Prob. with others, Table 5). While looking in the collection of all genomes with at least one Mn oxidase enzyme, MopA, MofA and BaMnxG were found alone with probabilities of 0.81, 0.831 and 0.691, respectively. The other two enzymes, McoA and PpMnxG, were rarely found alone (probability solo of 0.056 and 0.249, respectively). This pattern remains within subgroups of the greater population, with a few exceptions. In the gammaproteobacteria, MopA is more often found with other enzymes (0.416 vs 0.19 in the total population). In Deltaproteobacteria, McoA is almost always found solo (0.75) while BaMnxG is almost always found with other enzymes (0.906).

To determine which enzymes were most likely to be found together, we examined co-occurrence of Mn oxidation genes within genomes. The Pearson Chi square test is conducted to measure the strength of correlation between every bigram (pair) of genes. Four setups are considered in our study. First the correlations within the entire data set were studied, then we examined the co-occurrence patterns within two subsets of the dataset: proteobacteria, and gamma-proteobacteria.

To calculate the chi square we first built the contingency table as shown in Table 8. In the first row, we have the co-occurrences of the bigram of genes (gene 1 and gene 2) and the co-occurrences of gene 1 with all the other genes than gene 2 (the cases where gene 1 is alone are not considered, ¬ means not in logic). In the second row, we have all the co-occurrences of gene 2 with all the other genes than gene 1 and all the co-occurrences of the other pairs of genes than gene 1 and gene 2. To understand the chi square results, we need to examine some contingency tables. As seen in Table 9, McoA co-occurs with BaMnxG only once, while it co-occurs more frequently with other genes. BaMnxG co-occurs six times with other genes and McoA co-occurs 407 times with other genes. This is why the co-occurrences of this bigram of genes are not significant. As seen in Table 10, McoA and MopA co-occur 181 times. MopA co-occurs with other genes only 153 (less than with McoA, which is a strong indication). While McoA co-occurs 227 times with all the other genes (a little bit more than with MopA). That is why this co-occurrence of these two genes is significant. As seen in Table 11, MopA and PpMnxG co-occur 144 times. MopA co-occurs only 190 times with all other genes (a little bit more than with PpMnxG). On the other hand, PpMnxG co-occurs 229 times with all the other genes (a little bit more than with MopA). That is why this co-occurrence of these two is significant. Given that the co-occurrences of MopA and PpMnxG are not larger than with the other genes respectively, this pair has a lower effect size than with MopA-McoA and McoA-PpMnxG.

**Table 8.** Contingency table layout

|  | Gene 2 | ¬Gene 2 |
| --- | --- | --- |
| Gene 1 |  |  |
| ¬Gene 1 |  |  |

**Table 9.**
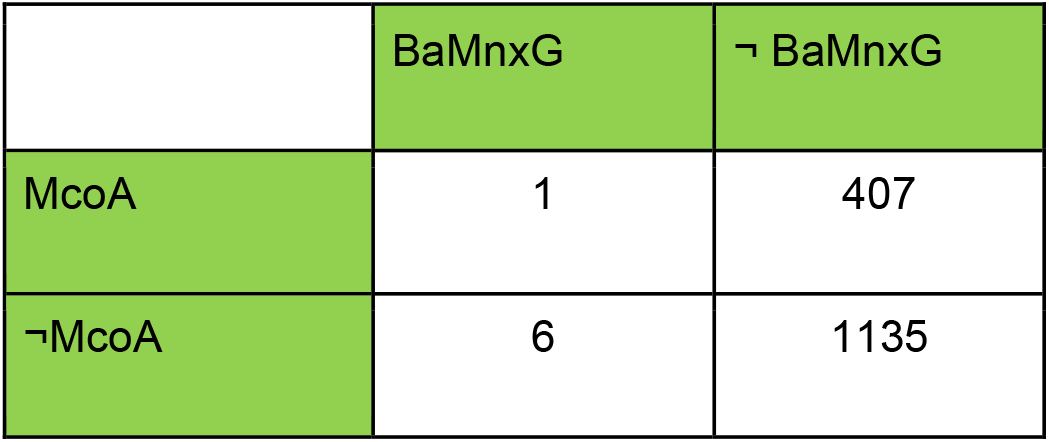
McoA and BaMnxG contingency table

|  | BaMnxG | ¬ BaMnxG |
| --- | --- | --- |
| McoA | 1 | 407 |
| ¬McoA | 6 | 1135 |

**Table 10.**
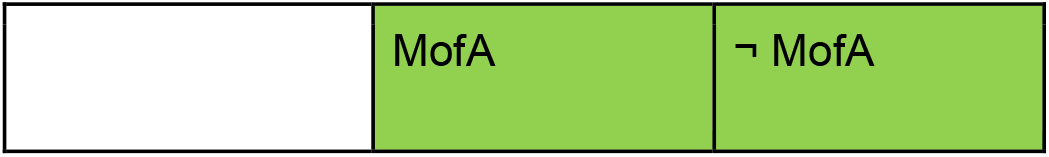
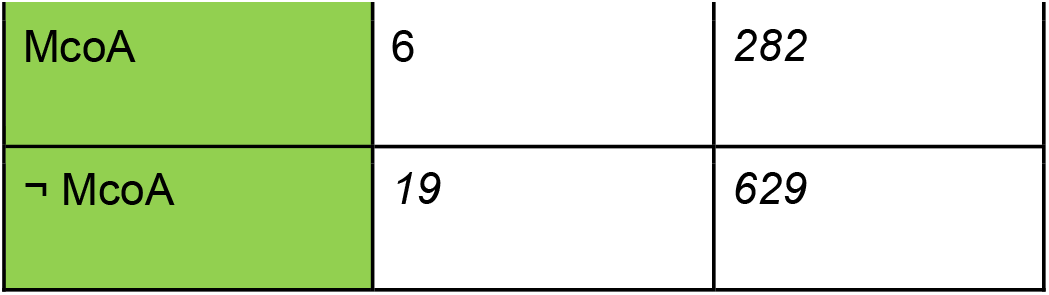
McoA and MofA Contingency Table

**Table 11.** MopA and McoA Contingency table

|  |  |  |
| --- | --- | --- |
|  | McoA | ¬ McoA |
| MopA | 181 | 52 |
| ¬ MopA | 107 | 452 |

Table 12 provides the chi square tests of the co-occurrence patterns of all the considered manganese oxidation genes. As we can see in this table, some bigrams of genes have significant chi square values with a medium effect size (Cramer’s V), like McoA-PpMnxG, McoA-MopA and MopA-PpMnxG while others have a significant chi square and a small effect size like PpMnxG-BaMnxG, McoA-BaMnxG and MopA-MofA. A third group did not show a statistically significant correlation like McoA-MofA and MofA-PpMnxG. When we examine co-occurrence of genes within the phylum proteobacteria (Table 13), and the class gamma-proteobacteria (Table 14), the most significant pairs with the highest effect sizes remain the same (McoA-PpMnxG; McoA-MopA and MopA-PpMnxG).

**Table 12.**
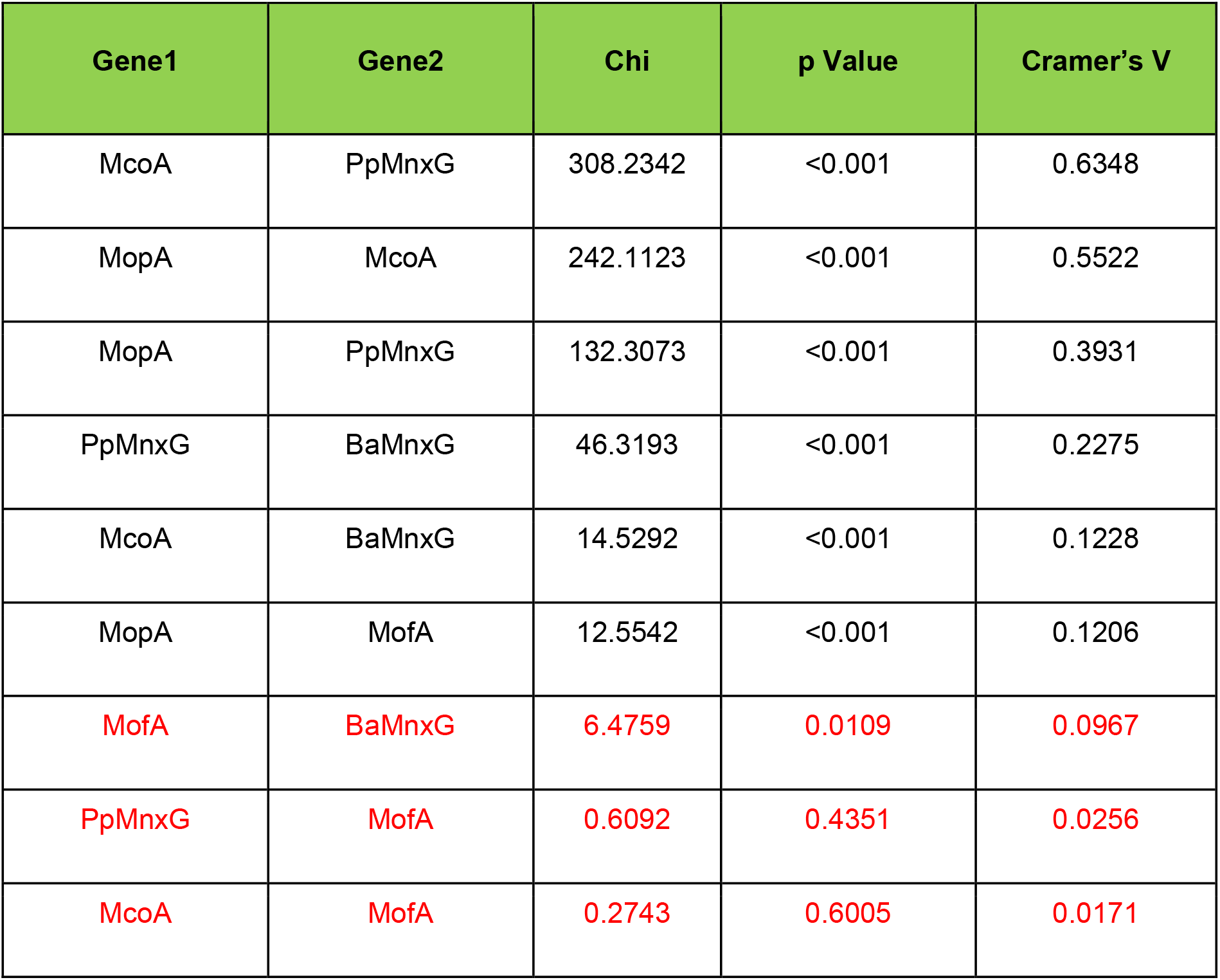
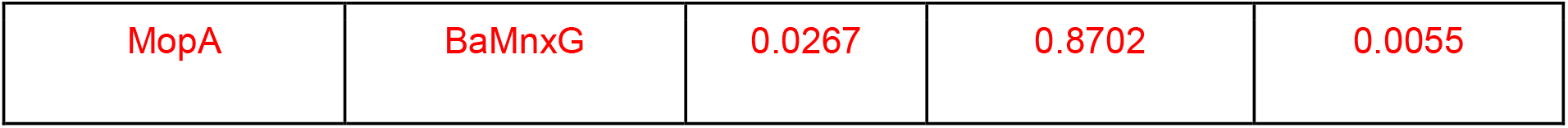
Gene-gene chi square in **all the data** Black text: bigrams with significant Chi values; red text: bigrams with insignificant Chi values.

| Gene1 | Gene2 | Chi | p Value | Cramer's V |
| --- | --- | --- | --- | --- |
| McoA | PpMnxG | 308.2342 | <0.001 | 0.6348 |
| MopA | McoA | 242.1123 | <0.001 | 0.5522 |
| MopA | PpMnxG | 132.3073 | <0.001 | 0.3931 |
| PpMnxG | BaMnxG | 46.3193 | <0.001 | 0.2275 |
| McoA | BaMnxG | 14.5292 | <0.001 | 0.1228 |
| MopA | MofA | 12.5542 | <0.001 | 0.1206 |
| MofA | BaMnxG | 6.4759 | 0.0109 | 0.0967 |
| PpMnxG | MofA | 0.6092 | 0.4351 | 0.0256 |
| McoA | MofA | 0.2743 | 0.6005 | 0.0171 |
| MopA | BaMnxG | 0.0267 | 0.8702 | 0.0055 |

**Table 13.** Gene-gene chi square in **proteobacteria**. Black text: bigrams with significant Chi values; red text: bigrams with insignificant Chi values.

| Gene1 | Gene2 | Chi | p Value | Cramer's V |
| --- | --- | --- | --- | --- |
| McoA | PpMnxG | 300.8431 | <0.001 | 0.6346 |
| MopA | McoA | 257.2006 | <0.001 | 0.5795 |
| MopA | PpMnxG | 143.9875 | <0.001 | 0.4175 |
| PpMnxG | BaMnxG | 58.2505 | <0.001 | 0.2583 |
| McoA | BaMnxG | 11.4397 | <0.001 | 0.1104 |
| MofA | BaMnxG | 7.7944 | 0.0052 | 0.1084 |
| PpMnxG | MofA | 3.845 | 0.0499 | 0.0653 |
| MopA | MofA | 2.2592 | 0.1328 | 0.0518 |
| MopA | BaMnxG | 0.7893 | 0.3743 | 0.0303 |
| McoA | MofA | 0.0083 | 0.9274 | 0.003 |

**Table 14.** Gene-gene chi square in gamma-proteobacteria

| Gene1 | Gene2 | chi2 | pValue | Cramers' V |
| --- | --- | --- | --- | --- |
| McoA | PpMnxG | 384.1001 | <0.001 | 0.8553 |
| MopA | McoA | 229.1318 | <0.001 | 0.6169 |
| MopA | PpMnxG | 170.682 | <0.001 | 0.5243 |
| MofA | BaMnxG | 19.7537 | <0.001 | 0.2002 |
| PpMnxG | BaMnxG | 3.6336 | 0.0566 | 0.0713 |
| PpMnxG | MofA | 1.7012 | 0.1921 | 0.0488 |
| McoA | MofA | 1.5578 | 0.212 | 0.0462 |
| McoA | BaMnxG | 0.3747 | 0.5405 | 0.0226 |
| MopA | BaMnxG | 0.0729 | 0.7872 | 0.0107 |
| MopA | MofA | 0.0037 | 0.9515 | 0.0024 |

### Probability of solo occurrence

The bigrams of genes shown above give an idea about the co-occurrence patterns between the genes but they don’t show the tendency of some genes to occur alone within a genome. To cover this aspect, we conducted another chi-square test. In this test, we used a contingency table like in Table 15, where we considered in the first row the number of the solo occurrences of a gene as opposed to its co-occurrences with other genes. In the second row, we considered the total number of solo occurrences of all the other genes as opposed to the number of their co-occurrences with other genes. As we can see in Table 16, all of the Mn oxidases, with the exception of BaMnxG, have a significant probability of being found solo, with the Cramer’s V effect size greatest for PpMnxG, McoA and MopA. With a probability cutoff of 0.05, BaMnxG is significantly likely to be found by itself in a genome. In the subset of proteobacteria genomes, the results are the same except now BaMnxG is significant (Table 17).

**Table 15.** Layout of the contingency table used to test if a gene occurs solo or in common with other genes

|  | Solo | Common |
| --- | --- | --- |
| Gene |  |  |
| ¬ Gene |  |  |

**Table 16.**
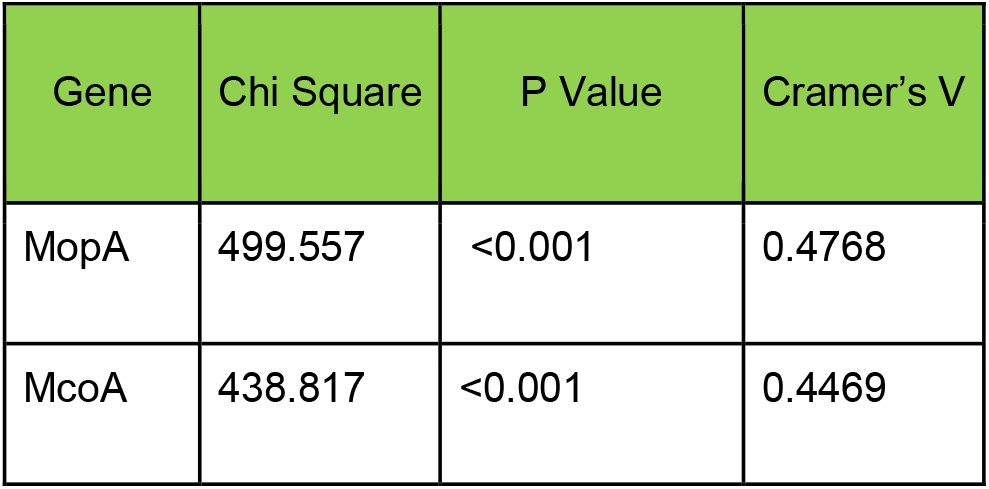
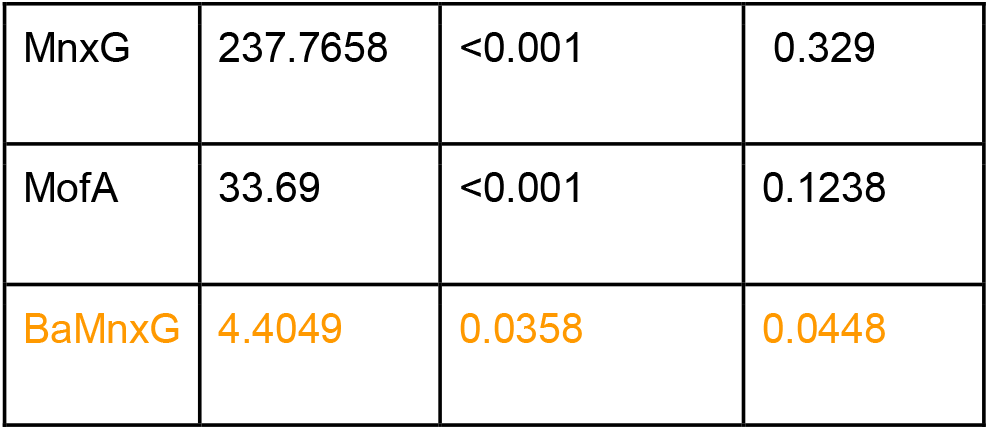
Chi square for solo occurrences of the genes within **the entire dataset**

**Table 17.** Chi square for solo occurrences of the genes **within proteobacteria**

| Gene | Chi Square | P Value | Cramer's V |
| --- | --- | --- | --- |
| MopA | 670.8704 | <0.001 | 0.5825 |
| McoA | 397.5534 | <0.001 | 0.4484 |
| MnxG | 200.5238 | <0.001 | 0.3185 |
| MofA | 14.8082 | <0.001 | 0.0865 |
| BaMnx<br>G | 25.9754 | <0.001 | 0.1146 |

## 4. Discussion

This work examined the distribution of putative Mn oxidase proteins in publicly available databases of protein and bacterial genome sequences; however, during the time it took to analyze these data, the number of sequences deposited in these databases grew considerably. Given the pace at which new genomes are sequenced, the results of this type of analysis will always be somewhat dated. The collection of sequenced genomes or proteins represents those organisms or proteins that have been deemed worthy of sequencing by researchers; therefore, the databases represent a biased subsample of the total diversity. While the number of sequences increased during the preparation of this paper, the biases that affect what gets sequenced likely have not changed.

Our results will also be affected by the sequences used to generate them. Our identification of Mn oxidase genes primarily in the phylum proteobacteria may be a result of using proteins identified in gamma- and alpha-proteobacteria as our queries. However, these were the proteins that have been identified as Mn oxidases; there likely are Mn oxidase enzymes that have not yet been characterized in other phyla. Repeating this project once more Mn oxidases have been identified will likely reveal different patterns. In addition, excluding the Mn oxidases (CopA, CotA, CueO, and MokA) with lower affinity for Mn^+2^ from our analysis likely altered the pattern. However, the need for high concentrations of Mn^+2^ to observe Mn oxidation suggests that these proteins are non-specifically oxidizing this metal and normally oxidize some other substrate. The higher affinity Mn oxidases are thus more likely to be true oxidases. An additional caveat to this work is the possibility that the homologs identified are not in fact Mn oxidases. For example, the heme-binding domain from the AHP MopA is found in other classes of proteins as well (20). Thus, identifying a potential Mn oxidase homolog in a genome is not sufficient to show that that organism oxidizes Mn. However, the overall pattern of which oxidases are found in which phyla should still be informative.

It has been proposed that the deltaproteobacteria and oligoflexia classes within the proteobacteria phylum be reclassified into four different phyla-level groups (Waite et al. 2020). Our data support the reclassification of deltaproteobacteria; the most common Mn oxidase proteins in this group are PpMnxG, MofA and BaMnxG while MofA and BaMnxG are relatively rare in the other proteobacteria (Table 5). There are not enough members of Oligoflexia with Mn oxidase homologs for us to draw any conclusions.

Our initial hypothesis was that more than one enzyme is required to complete the two-electron oxidation of Mn^+2^ to Mn^+4^. The MopA enzymes of *Erythrobacter* sp. SD-21 and *A. manganoxydans* SI85-9A1 perform a one electron oxidation to generate Mn^+3^ (18,36). Mn^+4^ oxides may be produced in these organisms via disproportionation or the action of a second enzyme, such as an MCO. For example, lignin-degrading white rot fungi have been shown to use a Mn peroxidase and an MCO together to oxidize lignin, with the Mn peroxidase producing Mn^+3^ (37,38). Therefore, we predicted that genomes would frequently have both an AHP and an MCO Mn oxidase gene present. However, four of the five Mn oxidases have a statistically significant probability of being solo (i.e. being the only Mn oxidase in the genome, Table 16). Even within the proteobacteria phylum, all five oxidases are found solo (Table 17) including the three oxidases that are found together in the genome of the model Mn-oxidizing bacterium *P. putida* GB-1. Therefore, our results do not support the original hypothesis. Indeed, biochemical analyses of Mnx complex of *Bacillus* sp. PL-12 have shown that this complex (in which BaMnxG is the presumed catalytic subunit) is capable of the two electron oxidation of Mn^+2^ to Mn^+4^ by itself (14).

The *Leptothrix discophora* sp. SS-1 genome encodes multiple potential Mn oxidase enzymes, including two *mofA* genes, two AHP genes similar to MopA, two genes predicted to encode an MCO similar to McoA and two homologs to BaMnxG (23). It is not clear how many of these proteins are capable of oxidizing Mn^+2^. Biochemical purification of the oxidase activity identified MofA as the Mn oxidase (39); however, deletion of this gene did not result in a loss of Mn oxidation (23). This raises the possibility that MofA is not a Mn oxidase; alternatively, it may be necessary to delete multiple Mn oxidation genes to lose oxidation activity, as has been observed in *P. putida* GB-1 (21,22).

The question remains, why do some species encode more than one Mn oxidase protein in its genome? As shown in Table 12 – 14, when genomes have more than one oxidase, the most common combination of proteins involve MnxG, McoA and MopA, the combination found in *P. putida* GB-1. Possibly this is an artifact of using the sequences of the *P. putida* GB-1 genes to search the databases. Nonetheless, different classes of bacteria exhibit different, characteristic patterns of Mn oxidase genes (Table 6). The firmicutes are dominated by one Mn oxidase, BaMnxG (Table 6) while the actinobacteria and proteobacteria have more equal distributions of several oxidases, as indicated by their higher Shannon diversity indices (Table 7). A possible explanation for this observation is that the different enzymes are optimized for oxidation under different conditions; organisms that grow or oxidize in a variety of environments therefore may use different oxidases depending on the growth conditions. In support of this hypothesis, various mutant strains of *P. putida* GB-1 exhibit different oxidation defects when the strains are grown on solid versus liquid media (22,40). A mutant strain lacking the gene for MnxG also behaves differently in normal versus low oxygen conditions (Geszvain, data not shown). In the firmicute *Bacillus* sp. SG-1 and related Bacillus species, the BaMnxG Mn oxidase is located in the spore coat and oxidation only occurs after sporulation (41,42). The more constricted conditions under which oxidation occurs in this phylum may be the reason why firmicutes often have only one oxidase enzyme in their genome.

## 5. Conclusion and Perspectives

By examining the distribution of Mn oxidase proteins in publicly available databases, we have shown that the genes for these proteins are predominantly found in the proteobacteria, firmicutes and actinobacteria genera, with each phylum having a characteristic distribution of Mn oxidases. Genomes that have a Mn oxidase protein frequently have just one. In genomes that do have more than one oxidase, MnxG McoA and MopA – the combination found in *P. putida* GB-1 – are often found together. Thus, while the situation found in *P. putida* GB-1 is not unique, it does not reflect a need for two enzymes to complete the oxidation of Mn. Instead, our results suggest that some organisms which need to oxidize Mn under a variety of conditions require oxidase enzymes optimized for those conditions. While the function of Mn oxidation in the cell remains unclear, these results further support the hypothesis that it plays a vital role in survival in the environment.

## 6. Methods

The amino acid sequences of selected MCO proteins are included in the supplemental data (Supplemental Figure S1). The AHP Mn oxidases (Figure S2) were screened for the presence of heme-binding domains using ScanProsite (https://prosite.expasy.org/scanprosite/). Using this program, *P. putida* GB-1 MopA was predicted to have two heme-binding domains; the amino acid sequence of the amino-terminal domain (Figure S3) was used to search for MopA homologs. Protein homology information was initially collected using the NCBI protein BLAST program, searching the non-redundant protein sequences (nr) database in late 2017. The nr database includes all non-redundant GenBank CDS translations as well as protein sequences from PDB, SwissProt, PIR and PRF excluding environmental samples from WGS projects. This resulted in a list of organisms that contained at least one putative Mn oxidase homolog. Representative genomes from these initial BLAST searches were used to select genomes for protein BLAST searches among completed genomes in IMG. All BLAST searches were limited to an E-value of 1 × 10^−50^, with the max target sequences set to the highest possible value and low complexity regions left unfiltered. Genome identifiers were used to label genomes that contained homologs of multiple proteins, with the presence or absence of a protein in a genome denoted as a binary variable. Taxonomic information for phyla and class was included for each genome and later used to calculate gene correlations to taxonomic groups. Over-representation of members within a single genus was limited by selecting five members of each genus to keep, using random selection via a randomization function in excel.

The statistical analysis is based on different techniques. First, we used co-occurrence tables of gene bigrams, a technique widely used in Statistical Natural Language Processing, which is a branch of Artificial Intelligence (46-47). To test our hypothesis, chi square tests were conducted with Cramer’s V to measure the effect size. Shannon diversity indices were also used to provide an orthogonal view to the one provided by the co-occurrence table.

## List of abbreviations

Mn: manganese
BaMnxG: the MnxG Mn oxidase found in *Bacillus* sp.
PpMnxG: the MnxG Mn oxidase found in *Pseudomonas putida* GB-1
MnOB: Mn-oxidizing bacteria
AHP: animal heme peroxidase
MCO: multicopper oxidase

